# Do we still need Illumina sequencing data?: Evaluating Oxford Nanopore Technologies R10.4.1 flow cells and v14 library prep kits for Gram negative bacteria whole genome assemblies

**DOI:** 10.1101/2023.09.25.559359

**Authors:** Nicole Lerminiaux, Ken Fakharuddin, Michael R. Mulvey, Laura Mataseje

**Author notes:** Correspondence: Laura Mataseje < >.

## Abstract

The best whole genome assemblies are currently built from a combination of highly accurate short-read sequencing data and long-read sequencing data that can bridge repetitive and problematic regions. Oxford Nanopore Technologies (ONT) produce long-read sequencing platforms and they are continually improving their technology to obtain higher-quality read data that is approaching the quality obtained from short-read platforms such as Illumina. As these innovations continue, we were interested in evaluating how much ONT read coverage produced by the Rapid Barcoding Kit v14 (SQK-RBK114) is necessary to generate high-quality hybrid and long-read-only genome assemblies for a panel of carbapenemase-producing *Enterobacterales* bacterial isolates. We found that 30X long-read coverage is sufficient if Illumina data is available, and that 100X long-read coverage is recommended for long-read-only assemblies. We found that Illumina polishing is still improving SNVs and INDELs in long-read-only assemblies. We also examined if antimicrobial resistance genes could be accurately identified in long-read-only data, and found that Flye assemblies regardless of ONT coverage detected > 94 % of resistance genes at 100% identity and length. Overall, the Rapid Barcoding Kit v14 and long-read-only assemblies can be an optimal sequencing strategy depending on the specific use case and resources available.

## Introduction

Genome assembly involves using genomic sequencing data (reads) to reconstruct an organism’s true genomic sequence^1^. There are currently two main types of sequencing data; (1) short-read sequencing platforms (e.g. Illumina) produce reads that are several hundred bases in length with high sequence accuracy, and (2) long-read sequencing platforms (e.g. Oxford Nanopore Technologies (ONT) and Pacific BioSciences) which produce reads that can be thousands of bases in length. Short-read sequencing data cannot resolve repeat regions larger than the library size^2^, which results in fragmented genomes and makes it difficult to determine whether genome fragments (contigs) are of chromosomal or plasmid origin. Long-read data spans more repeats which results in more contiguous assemblies, but has historically lagged behind short-read data in terms of error rates and insertions/deletions in homopolymer sequences^1,3–5^.

Genome contiguity becomes important when considering the mechanisms and drivers of antimicrobial resistance in bacterial pathogens. Plasmids are common vectors for horizontal gene transfer, and obtaining complete plasmid sequences is the most effective means to detect the coexistence of antimicrobial resistance genes on the same plasmid^6^, as well as the potential for that plasmid to transfer to other strains, species, or genera^7^. Long-read sequencing is usually essential to discern complete plasmid structures^6^, which often contain repetitive sequences that are difficult to resolve with short-read sequencing data alone^8^. Bacterial genome assemblies have been preferentially constructed with short-read-only data but are shifting to using short reads to polish long-read assemblies^9,10^.

ONT sequencing platforms are popular in microbial genomics due to their low cost in generating long-read data, their fast turnaround times, and their suitability for resource-limited environments^11^. ONT has two predominant workflows for whole genome sequencing: Rapid and Ligation. Rapid-based workflows result in longer read lengths and are quick to perform which makes it preferable for time-sensitive analyses (e.g. outbreak investigation), whereas Ligation workflows require more hands-on time but result in greater data yield^12,13^. Rapid-based workflows have been recommended for small plasmid recovery as small plasmids do not get fragmented in the Ligation-based workflows and are subsequently unavailable for sequencing^14^. ONT is continually pushing the boundaries of technological innovations and advancements are occurring at a rapid pace^15,16^. Many of the recent efforts with new sequencing kits has been to improve the read quality (Q-scores) through updates in the basecalling approach along with improvements in their library preparation and flow cell chemistry. The pores in the R9.4.1 flowcells (released in 2017) were made longer in the new R10.4.1 flow cells (released in 2022) which results in improved homopolymer sequencing^9^. In addition, error rates for newer chemistries have improved significantly, from ∼ 6 % for R9.4.1^3^ to ∼1 % for R10.4.1^9^.

Given the continual advances in ONT long-read sequencing, the need for additional short-read data for genome polishing of long-read assemblies is called into question. Several studies have examined bacterial genome assembly quality using the Ligation Sequencing Kit v14 (LSKv14)^6,17–19^, but no studies to date have examined assembly quality using the Rapid Barcoding Kit v14 (RBKv14). The objective of this project was to determine how much ONT read coverage produced by the Rapid Barcoding Kit v14 is necessary to generate high-quality hybrid and long-read-only genome assemblies. We used a dataset of 12 diverse carbapenemaseproducing Gram negative bacteria for validation, as these organisms are of clinical interest and contain a diverse plasmid population along with diverse antimicrobial resistance genes. We were also interested in comparing the ONT read quality produced by the Rapid Barcoding Kit v10 (SQK-RBK004), RBKv14 (SQK-RBK114-24), and LSKv14 (SQK-LSK114). Finally, we investigated if we could accurately identify the correct variant of acquired antimicrobial resistance genes in ONT-only assembled genomes without Illumina short-read data. These results provide guidance on when short-read data is beneficial for whole genome assemblies and when long-read data alone is sufficient.

## Results

We sequenced twelve isolates using three different methods: (1) Rapid Barcoding Kit v10 (RBKv10) loaded onto a R9.4.1. flow cell, (2) Rapid Barcoding Kit v14 (RBKv14) loaded onto a R10.4.1 flow cell, and (3) Ligation Sequencing Kit v14 (LSKv14) loaded onto a R10.4.1 flow cell. The average depth of coverage was 97X, 157X, and 77X for RBKv10, RBKv14, and LSKv14, respectively.

### How does read quality compare between v10 and v14 library preparation kits?

Both mean and median read quality scores (Q-scores) were improved for RBKv14 and LSKv14 relative to RBKv10, with LSKv14 producing the best Q-scores overall (Figure 1). The mean and median read length were similar between all read sets, but the read length N50 was best in LSKv14. These results confirm that the newer v14 library preparation kits result in higherquality reads relative to the older v10 kits.

**Figure 1:**
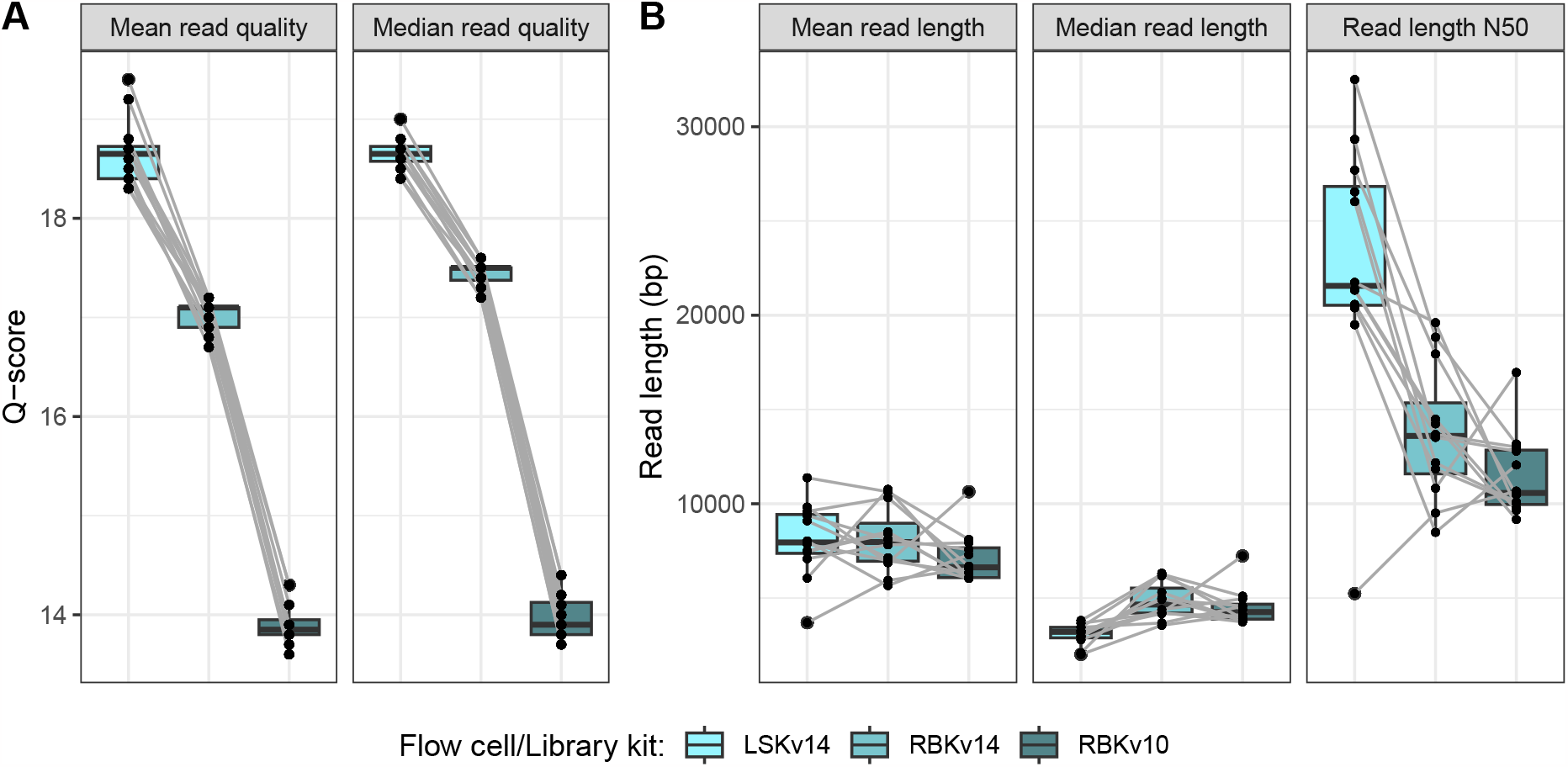
Mean and median read Q-score for Rapid Barcoding Kit v14, Ligation Sequencing Kit v14, and Rapid Barcoding Kit v10. Points represent isolates, and lines link the same isolates between different kits. A R9.4.1 flow cell was used for RBKv10, and R10.4.1 flow cells were used for RBKv14 and LSKv14.

We obtained average duplex read rates of 0.33 % and 5.6 % across all samples for the RBKv14 dataset and the LSKv14 dataset, respectively. Given the low rates of duplex reads in both RBKv14 and LSKv14 datasets generated in our hands combined with the extra computation time required for running multiple rounds of Guppy for duplex calls, we excluded the duplex tools process from our pipeline and used only simplex reads going forward.

While examining the read quality reports produced by NanoPlot, we noticed that the RBKv14 read sets had a more even read length distribution than the LSKv14 read sets. Despite the mean and median read lengths being similar, the LSKv14 reads tended to have a long, shallow tail for larger read lengths. An example for a single isolate is shown inFigure 2. Because of these results, and along with the shorter hands-on time required by the Rapid Barcoding Kit which is desirable for our specific diagnostic and outbreak workflows, the RBKv14 dataset was used for the remainder of the analyses.

**Figure 2:**
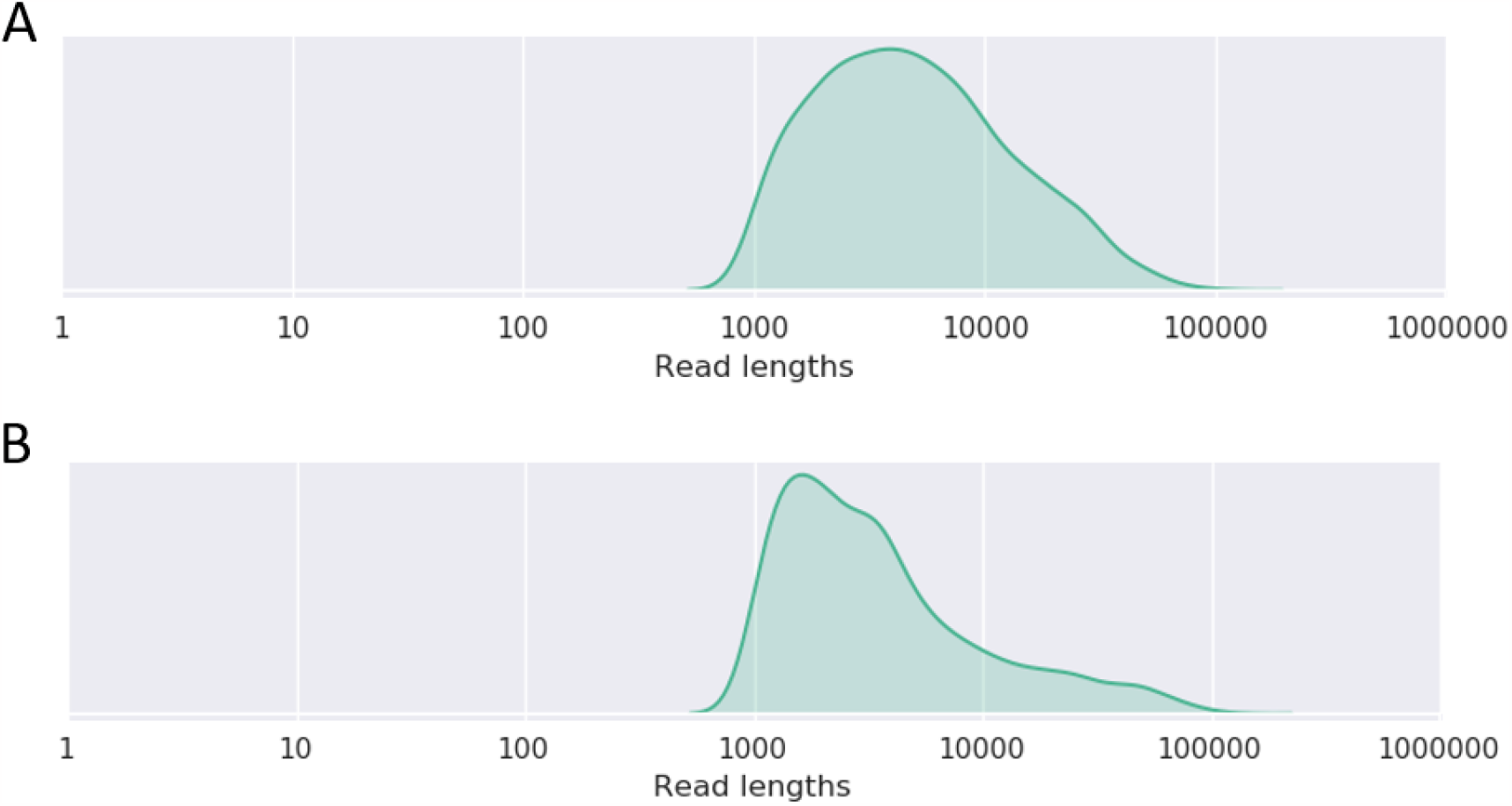
Histogram of read length distribution for isolateA sequenced to 100X average depth of coverage using the (A) Rapid Barcoding Kit v14 (RBKv14) and (B) Ligation Sequencing Kit v14 (LSKv14). Adapted from the NanoPlot “Read lengths vs Average read quality” plots. Read lengths represent bases.

### What depth of coverage (30X, 50X, 100X) is best for hybrid (Illumina & ONT) genome assembly?

Using the RBKv14 read sets from 12 isolates, we subsampled the reads to three different depths of coverage where achieved (30X (n=12), 50X (n=11), 100X (n=9)), assembled them using three different hybrid methods, and compared them to the reference genomes (see Methods). We examined assembly completeness or the presence of circular chromosome and plasmid replicons compared to the reference. Trends in assembly completeness appeared to be assembler-specific and generally did not change with increasing depth of coverage (Figure 3). Structurally, the Unicycler hybrid assemblies were the most similar to the reference genomes and typically contained small plasmids (< 10 kb) that were absent or duplicated in other assemblies. Flye frequently duplicated small plasmids (between 4.1 - 9.3 kb) or was missing small plasmid replicons altogether (between 1.4 - 5.2 kb). Raven also omitted plasmids (between 1.4 - 111.3 kb). Raven and Unicycler did not circularize the chromosome in several instances, but Raven could resolve those structures with additional sequencing depth. Illumina polishing did not change the number or completeness of replicons, indicating this graph looks identical for the Flye and Raven long-read-only assemblies.

**Figure 3:**
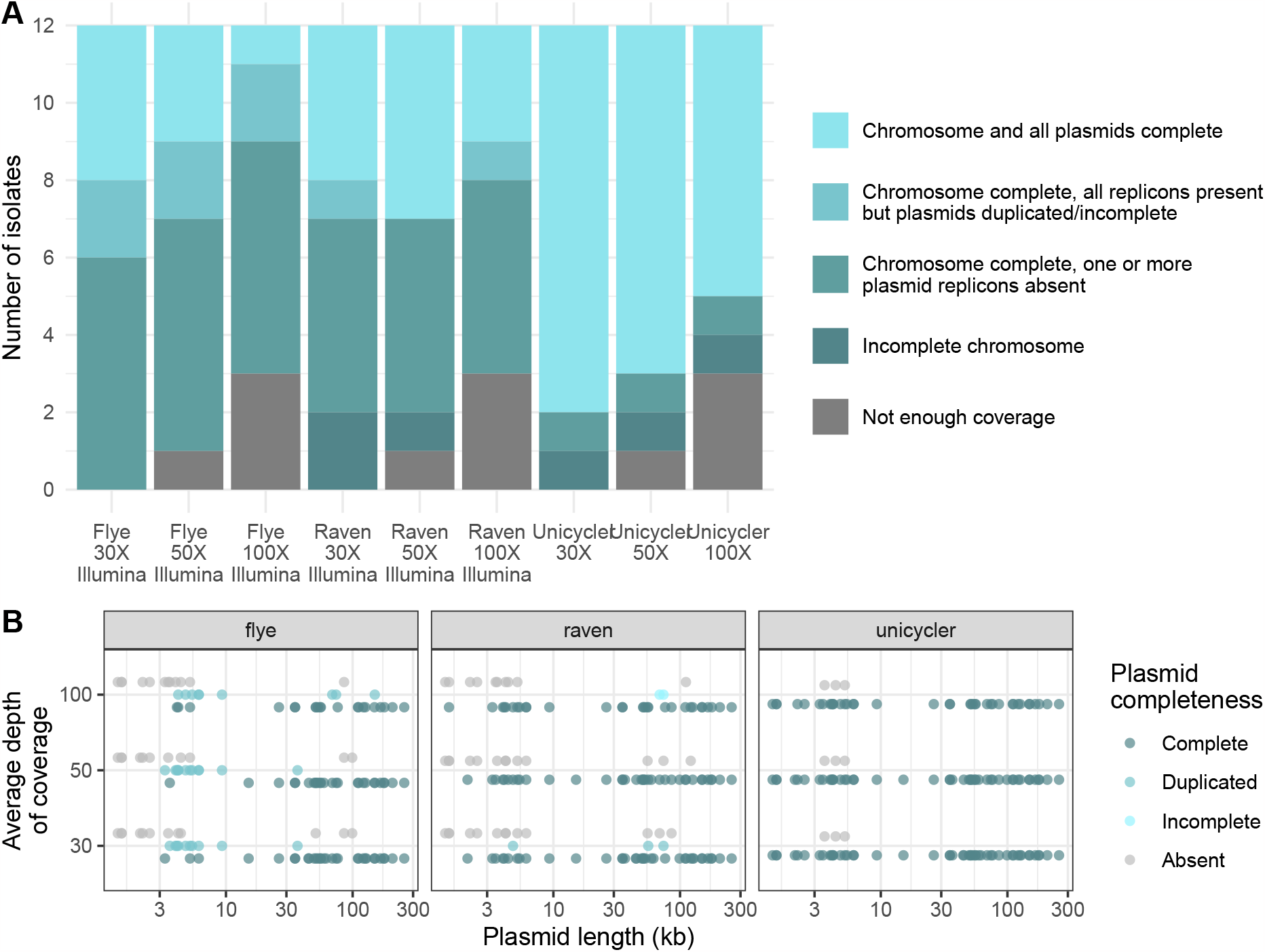
Hybrid assembly completeness relative to the reference genomes. (A) Completeness across all replicons. One isolate did not have enough coverage at 50X, and an additional two isolates did not have enough coverage at 100X. (B) Completeness of plasmids. Absent indicates the replicon was missing from the assembly.

We compared the hybrid assemblies to the reference genomes to determine what depth of ONT read coverage produces the best hybrid assemblies. We found that there was no difference between any metric for the read depths tested regardless of assembly method (Figure 4 A). When comparing to the reference genomes, the Unicycler assemblies had an average of 1.5 single nucleotide variants (SNVs) per 100 kb regardless of read depth and 0.30 (30X and 50X) or 0.29 (100X) small (< 60 bp) insertions/deletions (INDELs) per 100 kb. The Flye and Raven assemblies had fewer SNVs and INDELs than the Unicycler hybrid assemblies; Flye assemblies had an average of 0.15, 0.08, and 0.10 SNPs per 100 kb and 0.17, 0.07, and 0.09 INDELs per 100 kb for 30X, 50X, and 100X coverage, respectively. Raven assemblies had an average of 0.19, 0.18, and 0.05 SNPs and 0.30, 0.43, 0.27 INDELs per 100 kb for 30X, 50X and 100X coverage, respectively. The SNVs in the Flye and Raven assemblies were primarily located in transposases, resolvases, and intergenic regions, whereas the SNVs in the Unicycler assemblies were overwhelmingly found in rRNA and tRNA regions. Overall, this indicates an ONT read set with a minimum average depth of coverage of 30X is sufficient to produce a high-quality genome assembly if Illumina data is available.

**Figure 4:**
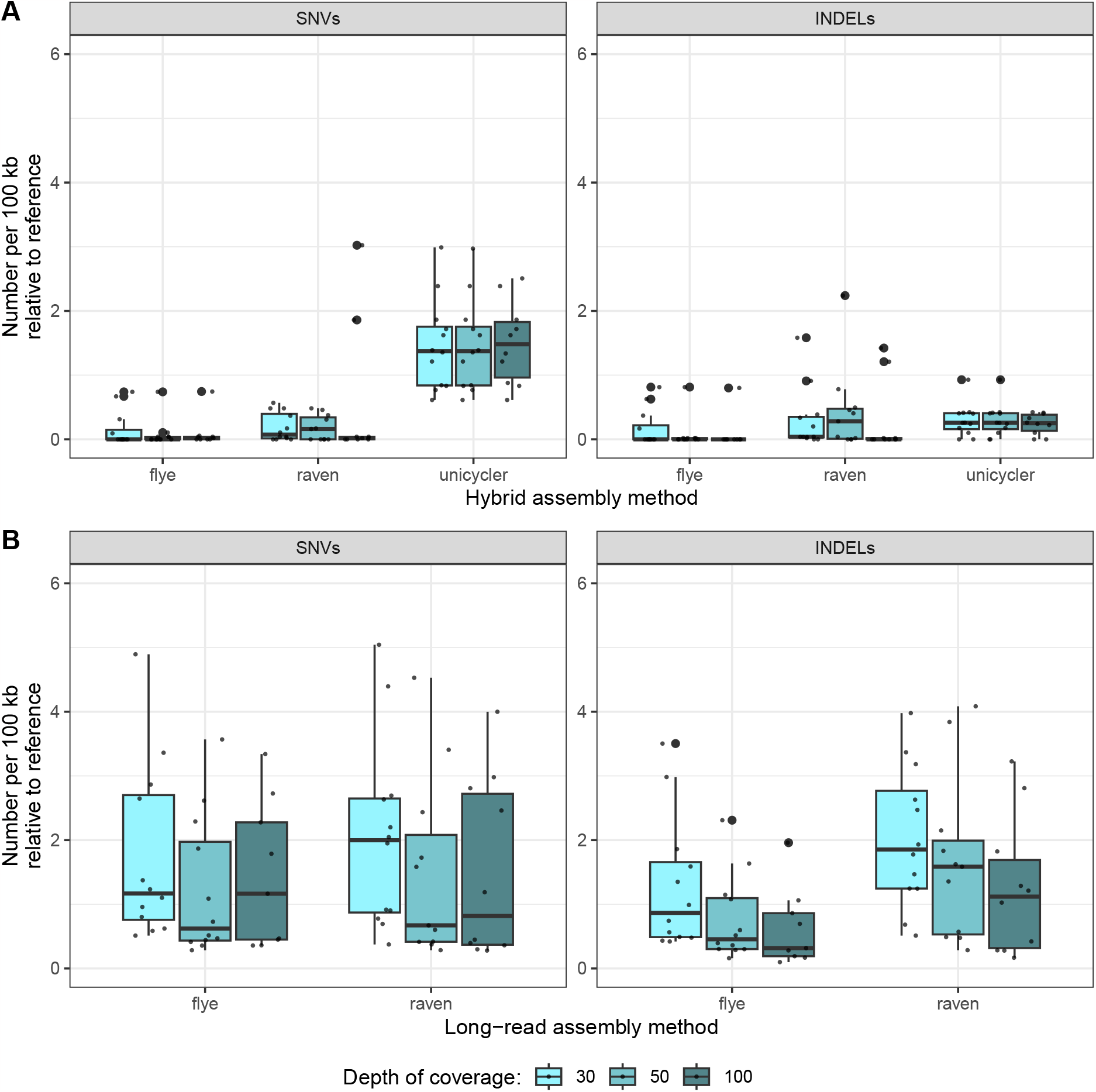
Comparison of single nucleotide variants (SNVs) and small insertions/deletions < 60 bases (INDELs) in (A) hybrid assembly methods and (B) long-read-only methods at three different average read depth of coverage to the reference genomes. Lines in the center of boxplots represent the mean.

### What depth of coverage (30X, 50X, 100X) is best for long-read-only (ONT) genome assembly?

To determine what depth of ONT coverage produces the best long-read-only assemblies, we assembled the 30X, 50X, and 100X RBKv14 read sets using two different long-read-only methods Flye and Raven (see Methods), then compared these to the reference genomes. We chose to not assess the long-read-only mode in Unicycler (a Miniasm^20^ wrapper) given poor performance in a recent study^11^. The total number of SNVs and INDELs improved with additional read depth, with 100X coverage producing the fewest number of assembly errors, followed by 50X (Figure 4 B). Results were comparable between Flye and Raven; addtional long-read coverage improved the average number of SNVs per 100 kb from 1.75 (Flye 30X) and 2.05 (Raven 30X) to 1.44 (Flye 100X) and 1.52 (Raven 100X), as well as the average number of INDELs per 100 kb from 1.28 (Flye 30X) and 2.04 (Raven 30X) to 0.63 (Flye 100X) and 1.25 (Raven 100X). If Illumina data is unavailable, higher depth of coverage for ONT read sets is preferable for more accurate genomes, with improvements in SNVs and small insertions/deletions seen up to 100X.

### How close are we to using ONT data to produce Illumina-quality assemblies?

To determine how much Illumina short-read polishing is improving the long-read only assemblies, we compared the assemblies before and after polishing (Figure 4). When Illumina data is used to polish Flye and Raven assemblies, the number of SNVs per genome is reduced from a mean of 77 (Flye) and 82 (Raven) SNVs to 5 or fewer SNVs at 100X coverage relative to the respective reference genomes. Similarly, Illumina polishing results in a reduction of INDELs per genome from a median of 33 (Flye) and 66 (Raven) to 0, with few exceptions. Overall, this indicates that Illumina data is still useful in resolving small-scale SNVs and INDELs in long-read-only assemblies.

### Can antimicrobial resistance genes be accurately identified in long-read-only (ONT) assemblies?

We compared the antimicrobial resistance gene profiles from the long-read-only assemblies to the reference genomes to determine if resistance genes could be accurately identified (100 % identity over the full gene length) without Illumina data. A total of 186 antimicrobial resistance genes (including duplicates within isolates) were detected in the reference genomes, representing 69 unique antimicrobial resistance genes, and the number per isolate ranged from 2 to 29 (mean: 15.5). Flye assemblies at all depths and Raven at 100X accurately identified the largest proportion of antimicrobial resistance genes (> 90%) (Table 1), indicating additional read depth improves coverage of Raven assemblies but not Flye assemblies. The genes that were not correctly identified were either present with sequence errors relative to the reference, had duplicate copies that did not exist in the reference, or were absent altogether (Figure 5).

**Table 1:**
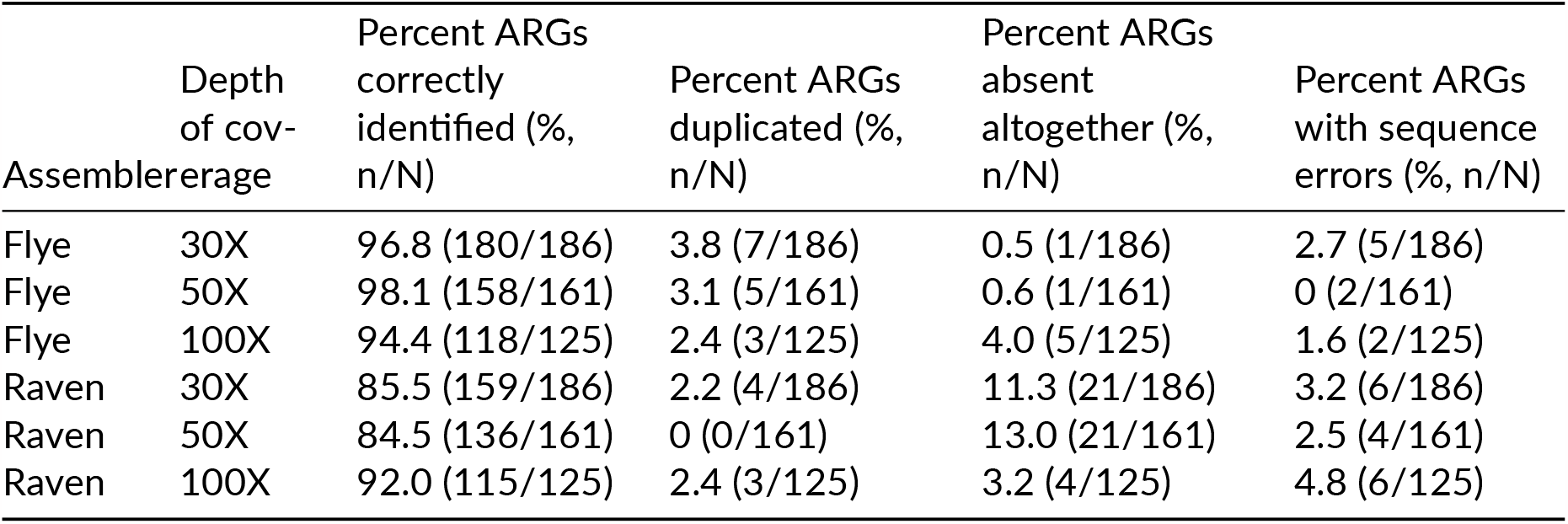
Proportion of correctly identified (100 % identity and length) antimicrobial resistance genes (ARGs) in long-read-only assemblies.

**Figure 5:**
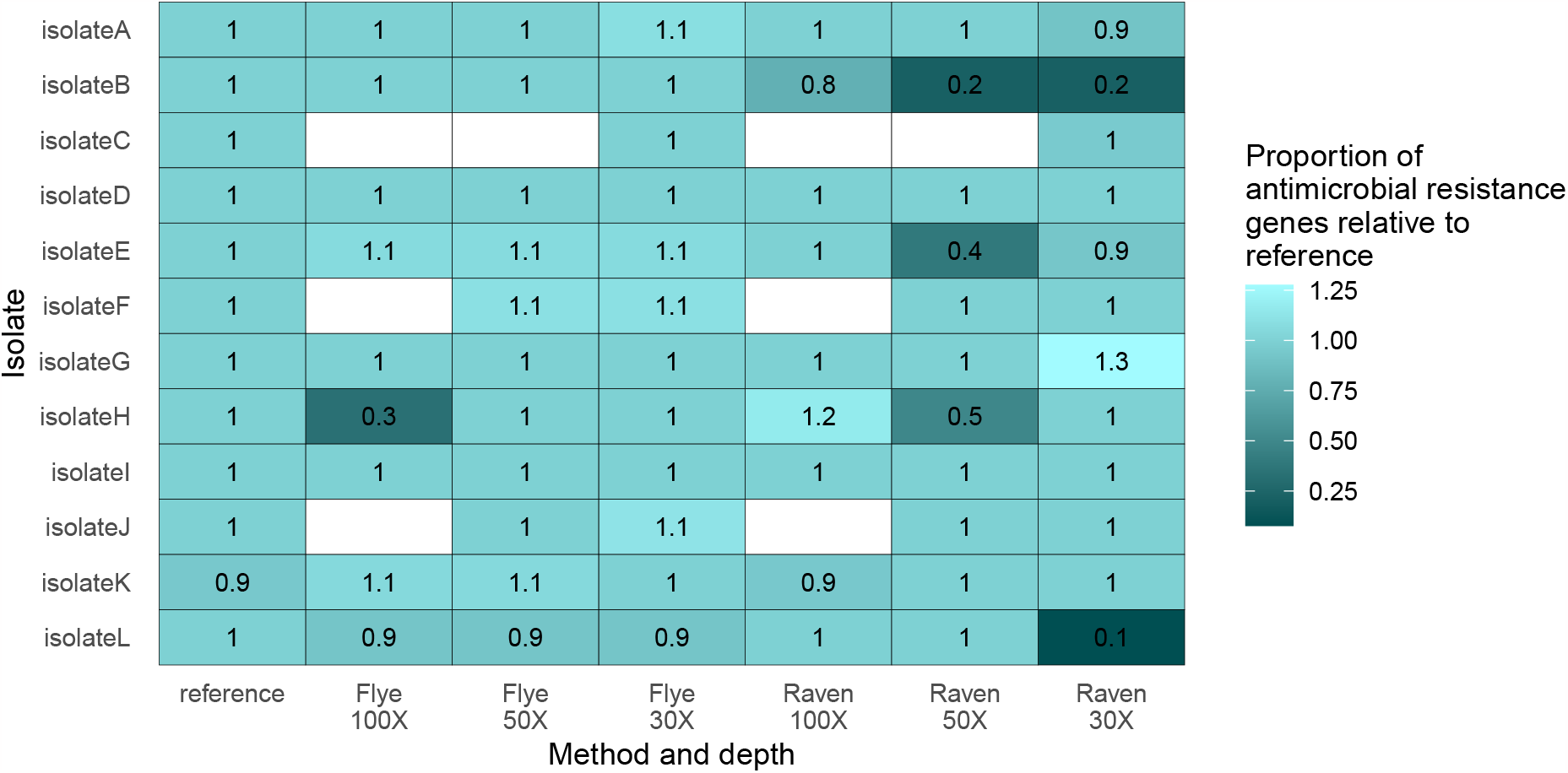
Proportion of correctly identified antimicrobial resistance genes across long-read-only methods relative to the reference genomes. Missing values indicate isolates that did not have enough read depth. Proportions below one indicate genes are missing, and proportions above one indicate genes were duplicated. The proportion value below in 1 the isolateK reference and Raven 100X corresponds to *bla*_OXA-232_ that was present but split over the start/end of a 6.1 kb plasmid.

We found that 3/12 isolates had 100% agreement to the reference antimicrobial resistance gene sequences and profiles over all assembly methods and depths. Flye was more consistent in correctly identifying antimicrobial resistance genes than Raven, wherein only isolateH had a large misassembly at Flye 100X resulting in four absent antimicrobial resistance genes. There were 3 isolates in which Flye duplicated either a 6.1 kb *bla*_OXA-232_ plasmid or a 36.6 kb *bla*_KPC-3_ plasmid, resulting in extra resistance gene copies relative to the reference. In isolateL, Flye could not resolve a 85 kb plasmid with a 10 kb *bla*_KPC-3_-encoding duplication, resulting in a missed copy of *bla*_KPC-3_. For the Raven assemblies, there were 4 isolates in which one or more assemblies omitted a large plasmid (> 69 kb), resulting in missed antimicrobial resistance genes. There were also two instances of Raven duplicating large plasmids (56 and 74 kb), resulting in extra resistance gene copies. The proportion value below 1 for the isolateK reference and Raven 100X corresponds to *bla*_OXA-232_ that was present but split over the start/end of a 6.1 kb plasmid. There were several instances (2.6%, 25/944 total genes in long-read-only assemblies) where the antimicrobial resistance gene was present but contained single nucleotide variants (SNVs) relative to the reference genomes (Table 1). Genes with errors common to multiple long-read assemblies were *tetB, bla*_TEM-1B_, *aadA* variants, *ermB*, and *mph(A)*, and most errors 72.0%, 18/25) were corrected with Illumina polishing.

## Discussion

We constructed assemblies with different ONT read depths to determine if additional ONT read depth improves long-read-only assemblies or hybrid assemblies with Illumina short reads. We found that having additional RBKv14 long-read depth of coverage did not improve assembly quality when Illumina data was available, but additional read depth did improve SNVs and INDELs in long-read-only assemblies. Illumina short-reads are still improving SNVs and INDELs in long-read-only assemblies (>75 SNVs, >29 INDELs per 5.2 Mb genome). We examined the RBKv14 long-read-only assemblies for antimicrobial resistance genes, and found that additional read depth improved gene detection in Raven assemblies, but not for Flye assemblies. All Flye assemblies regardless of ONT coverage detected > 94 % of resistance genes; those that were missed were completely absent from the genome due plasmid replicon misassemblies and minor SNV errors. As mentioned by^18^, the optimal sequencing and assembly strategy will depend on the specific use case and resources available, and that strategy will likely evolve rapidly over the next several years as the technology continues to improve.

If Illumina data is unavailable, we found that more long-read coverage (up to 100X) resulted in better assemblies with fewer SNVs and INDELs. Other LSKv12 and LSKv14 studies have found the minimum coverage for assembling SUP long-read-only data without substantial improvements was 20X^18^, 30X^21^, or 40X^9^.^9^ found 40X coverage of long-read only resulted in < 4 INDELs per 100 kb but did not examine SNVs. Similar to our results,^18^ found the number of SNVs did not greatly improve with more than 20X coverage, but additional coverage appeared to further to reduce SNVs and INDELs at different magnitudes in different species.

If Illumina data is available, minimal ONT long-read coverage is needed (30X was sufficient) to obtain high-quality genomes. Others have found as little as 10X coverage or lower is sufficient for accurate reconstruction of chromosomes and plasmids with Unicycler hybrid with LSKv12 on R10.3 flow cells^18^ and with LSKv10^22^. This makes sense given the approach used by Unicycler, wherein the long reads are only used as bridges to connect short-read contigs^22^. In our study, Illumina short-read data is still improving SNVs and small insertions/deletions in Flye and Raven RBKv14 assemblies, so Illumina data is recommended if the genomes are to be used in high-resolution downstream analyses such as SNP analysis to track transmission events.

We chose Flye and Raven as our long-read assemblers, both of which performed well relative to other long-read assemblers^1^. Small plasmids are known to be underrepresented in long-read-only assemblies either due to the assembly strategy or the ONT library preparation protocol^14,22^; however, even using an ONT Rapid kit which is recommended over Ligation kits for assembling small plasmids^14^, both assemblers still struggled with small plasmid assembly, which resulted in only partially complete assemblies for most isolates (Figure 3). Flye duplication of small plasmids has been observed in other studies^1,18,23^ and did not seem impacted by average depth of coverage in this study. Raven was better at resolving smaller plasmids than Flye but would occasionally omit a larger plasmid from the assembly altogether. This underrepresentation of small plasmids was not observed in the Unicycler assemblies. If small plasmids are important, we recommend including Illumina data to accurately identify them. If large plasmids are important, we recommend using Flye to assemble.

Long-read-only LSKv14 assemblies have been shown to be accurate for detecting acquired antimicrobial resistance genes, virulence factors, and performing cgMLST analysis^17,19^. Equivalent data for RBKv14 has not yet been published, but^21^ tested various genera with RBKv10 on R10.3 but found Illumina data resulted in better AMR gene detection and cgMLST, and both the library preparation kit and flow cells have newer versions. Here, we assessed whether ONT-only RBKv14 assemblies were sufficient to accurately identify acquired antimicrobial resistance genes. There was several instances where the resistance gene did not match the reference genomes due to single nucleotide variants, and several of these variants remained after Illumina polishing. We found that Flye accurately identified all resistance genes in most isolates, but incorrectly overestimated the number of copies of certain genes due to duplication of smaller plasmids (< 10 kb). In contrast, Raven performed better with small plasmids but would occasionally omit a large replicon (> 69 kb) containing antimicrobial resistance genes. The smallest plasmid containing an antimicrobial resistance gene in our dataset was the 6.1 kb ColE replicon encoding *bla*_OXA-232_ found in two isolates; this plasmid was duplicated but present in all Flye assemblies and complete in the Raven 100X assemblies but missing at the 30X and 50X assemblies for one isolate. If small plasmids carrying antimicrobial resistance genes are of interest, we recommend using RBKv14^14^ and comparing small plasmid content assembled with Raven and hybrid Unicycler if Illumina data is available. If general antimicrobial resistance gene detection across all replicon sizes is the goal with long-read-only data, we recommend Flye over Raven but verification of plasmid duplication is recommended to resolve the number of repeated antimicrobial resistance genes if required.

We did not investigate the impact of different basecalling models (Fast, High-accuracy (HAC) and Super-accuracy (SUP)) on the assemblies generated, and instead opted to use the SUP model. If computing and time resources are available, others have recommended using the SUP model for basecalling to obtain the best results, and polishing with Medaka is worthwhile for improved performance regardless of long-read coverage^18,22,24^.

We investigated the difference in read quality metrics between RBKv10, RBKv14, and LSKv14 library preparation kits, and found that read Q-scores were highest with LSKv14 followed by RBKv14, but read length distribution was best with RBKv14. Both LSKv14 and RBKv14 had better read quality metrics than RBKv10, which agree with other studies^6,11,17,18^. While LSKv14 had higher Q-scores, the read length distribution for RBKv14 had a better representation of longer reads in the RBKv14 datasets, whereas the LSKv14 read size was skewed towards shorter reads, which has been observed previously with older kit/flowcell versions (SQK-LSK108 & SQK-RAD002 on R9.4.1 flow cells)^12,13^. We detected low proportions of duplex reads from RBKv14 (average 0.33 %) and LSKv14 (average 5.6 %); this agrees agree with^18^ who found the proportion of duplex reads with LSKv14 was < 7 % for each isolate. With further optimization, this proportion would hopefully increase. Long-read-only assemblies will likely improve with the exclusive use of duplex reads, but current protocols are not generating enough duplex reads and consequently the coverage required to make this feasible^18^.

## Conclusions

It is marvelous that genomes assembled with ONT-only data contain fewer than 100 SNVs relative to the gold-standard hybrid methods. In a 5.2 Mb genome, this represents an error rate of less than 0.002 % per base. ONT RBKv14 long-read data has shown it can accurately detect > 94% of antimicrobial resistance genes with as little as 30X coverage when Flye is used as the assembler. However, Illumina data is still improving SNVs and INDELs in SUP-basecalled RBKv14 data, indicating Illumina data is still recommended for high-resolution downstream analyses. As ONT continues to innovate and upgrade their technology, these numbers will no doubt improve to attain Illumina-quality assemblies using only long-read data within the next few years.

## Data availability

Sequencing reads are deposited in the SRA database under BioProject PRJNA1020811. BioSample and GenBank accessions can be found inTable 2. Analysis code is available upon request.

**Table 2:**
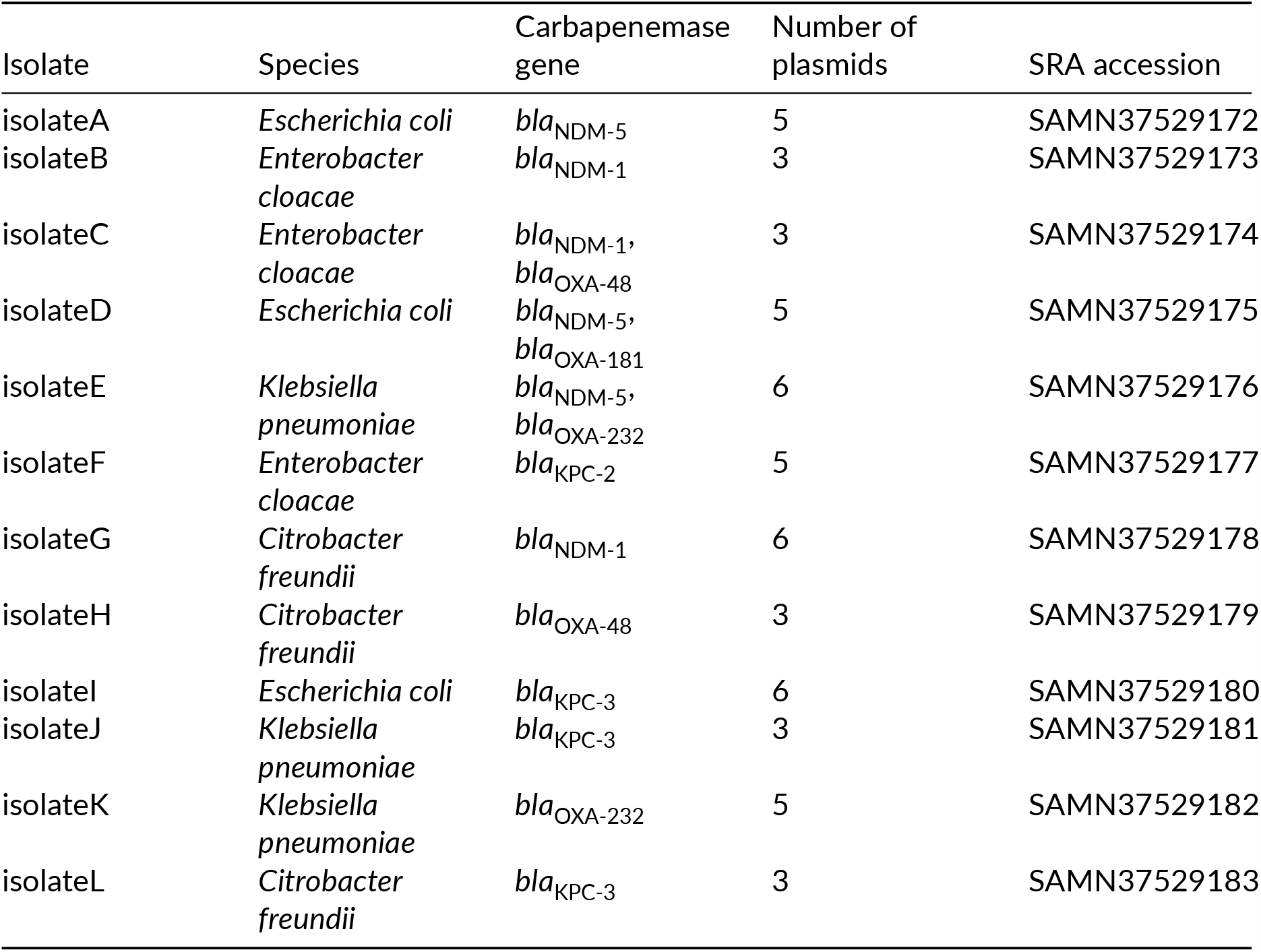
Panel of isolates.

## Methods

### Panel of Gram negative organisms

We selected a panel of twelve Gram negative carbapenemase-producing organisms that were collected as part of the Canadian Nosocomial Infection and Surveillance Program (CNISP)^25^. These twelve isolates were selected to represent various species, carbapenemase genes, and diverse plasmid content (Table 2).

### DNA extraction and sequencing

Cultures were streaked from freezer stocks, inoculated into LB broth culture, and grown overnight at 37° Celsius. Genomic DNA was extracted using the Epicentre MasterPure™ Complete kits (Mandel Scientific, Guelph, ON, Canada) following the manufacturer’s protocol for total DNA from cell samples. The same DNA extract was used for both short-read Illumina and long-read ONT sequencing where possible.

We performed three separate ONT library preparation protocols: (1) Rapid Barcoding Kit v10 (SQK-RBK004) loaded onto a R9.4.1 flow cell, (2) Rapid Barcoding Kit v14 (SQK-RBK114-24) loaded onto a R10.4.1 flow cell, (3) Ligation Sequencing Kit v14 (SQK-LSK114) with Native Barcoding Kit (EXP-NBD114) loaded onto a R10.4.1 flow cell. All twelve isolates were run concurrently on unused flow cells for each library preparation kit/flow cell combination specified. All sequencing was performed with the MinION Mk1B (ONT, Oxford, Oxfordshire, UK).

Short-read Illumina libraries were created with TruSeq Nano DNA HT sample preparation kits (Illumina, San Diego, CA, USA) following the manufacturer’s protocol. Paired-end, 301 bp indexed reads were generated on an Illumina MiSeq™ platform (Illumina).

### Read processing

ONT reads were basecalled and demultiplexed with Guppy v6.4.6 using the Super High Accuracy model (ONT). Adaptors and barcodes were trimmed with Porechop v0.2.3_seqan2.1.1 (https://github.com/rrwick/Porechop) and reads were filtered for Q-score > 8 and length > 1000 bases with Filtlong v0.2.1 (https://github.com/rrwick/Filtlong) (–min_length 1000 – min_mean_q 85). Illumina reads had adaptors trimmed and were filtered for an average Q-score > 30 with Trim Galore v0.6.7 (https://github.com/FelixKrueger/TrimGalore) (–paired –quality 30 –fastqc). FastQC v0.11.9 (https://github.com/s-andrews/FastQC) and NanoPlot v1.28.2^26^ were used to assess quality control metrics for Illumina and ONT reads respectively.

We searched for concatemeric reads in the ONT r10.4.1 datasets using duplex-tools v0.3.1 as recommended by ONT (https://github.com/nanoporetech/duplex-tools). Given the low rates of duplex reads in both RBKv14 and LSKv14 datasets generated in our hands with our standard workflows combined with the extra computation time required for running multiple rounds of Guppy for duplex calls, we excluded the duplex tools process from our pipeline and used only simplex reads going forward.

### Generating read subsets and assemblies

We generated assemblies to assess ONT assembly quality using five different methods at three different read depths: 30X, 50X, and 100X average depth of coverage, which resulted in 15 assemblies per isolate. Less than 100X coverage was obtained for 7/12, 3/12, and 10/12 isolates from RBKv10, RBKv14, and LSKv14, respectively. Of those, 1/12, 1/12, and 6/12 obtained less than 50X coverage for RBKv10, RBKv14, and LSKv14, respectively. Isolates that did not attain the target depth of coverage were excluded from that depth category. Reads were subsampled with seqtk v1.3 (https://github.com/lh3/seqtk) by multiplying the depth of coverage required by an average genome size of 5.2 Mb then dividing by the mean read length to obtain the number of reads required for the target depth. Each read subset was passed to five different assembly pipelines shown inFigure 6. The assembly workflows were managed using Snakemake^27^. Assemblies were generated using Unicycler v0.5.0^28^, Flye v2.9.2^29^ (–nano-hq), or Raven v1.8.1^30^ (–graphical-fragment-assembly). Assemblies were polished with long reads using Medaka v1.7.2 (https://github.com/nanoporetech/medaka) (model r1041_e82_400bps_sup_g615) and optionally with short reads using Polypolish v0.5.0^31^ and POLCA from MaSuRCA v4.0.9^32^ if specified. The five assembly methods are summarized below:

**Figure 6:**
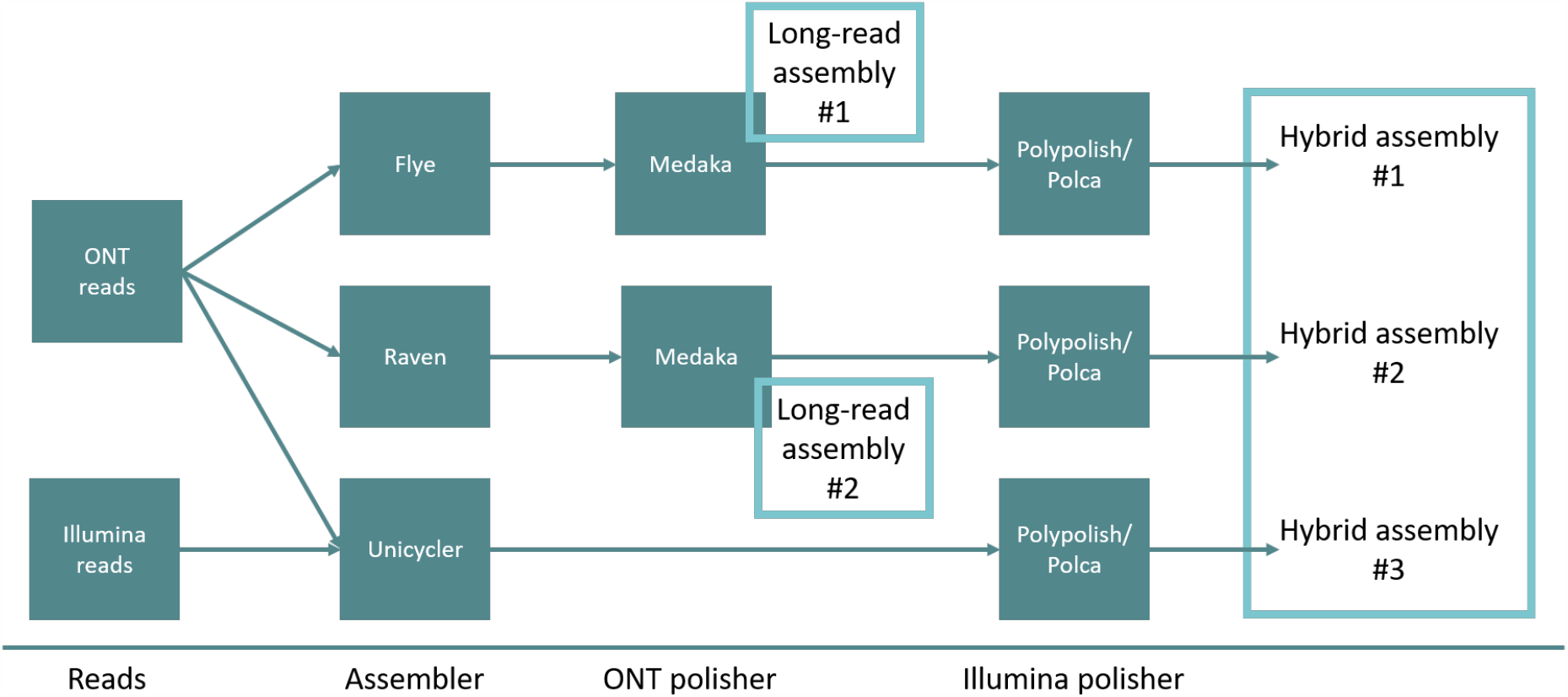
The five hybrid and long-read assembly methods compared

1. Hybrid assembly #1: Flye long-read assembly, Medaka long-read polishing, and Polypolish and POLCA short-read polishing
2. Hybrid assembly #2: Raven long-read assembly, Medaka long-read polishing, and Polypolish and POLCA short-read polishing
3. Hybrid assembly #3: Unicycler hybrid assembly, and Polypolish and POLCA short-read polishing
4. Long-read assembly #4: Flye long-read assembly, and Medaka long-read polishing
5. Long-read assembly #5: Raven long-read assembly, and Medaka long-read polishing

### Generating reference genomes

We used Trycycler v0.5.3^4^ to generate a set of reference genomes from the Rapid Barcoding Kit v14 (SQK-RBK114-24) run. Twelve read sets randomly subsampled representing an average depth of coverage of 60X using seqtk v1.3^33^. Four read sets each were assembled by Flye v2.9.2^29^, Raven v1.8.1^30^, and Miniasm v0.3_r179^20^. These assemblies were input into the Trycycler pipeline and the consensus assemblies were polished using both short-read and long-read polishers as described above. Three small plasmids (< 5 kb) found in Unicycler assemblies were absent from the Trycycler references, and these were manually added to the reference genomes (1.4 kb and 1.5 kb in isolateI, and 4.2 kb in isolateK).

### Comparing read quality, assembly quality, SNVs, and antimicrobial resistance genes

Read quality was assessed using NanoPlot v1.28.2^26^. Average depth of coverage was estimated by dividing the total bases after quality trimming by an average genome size of 5 Mb.

Each of the isolate genomes were aligned to the reference isolate genome using the dnadiff tool from MUMmer v3.23^34^ to obtain the number of single nucleotide polymorphisms (SNVs), small insertions/deletions < 60 bp (indels), large insertion/deletions > 60 bp, and size of all large insertions. The resulting positions in the .snps file were searched against genome annotations produced by Bakta v1.6.1^35^ to identify the locus of the SNP. StarAMR v0.9.1^36^ was used to identify antimicrobial resistance genes.

Plots were generated in R v4.3.0 (https://www.R-project.org/) using ggpubr v0.6.0 (https://cran.r-project.org/web/packages/ggpubr/index.html) and the tidyverse packages^37^.

## Acknowledgements

We would like to dedicate this work to the late Dr. Michael Mulvey, who’s vision for advancing genomics for the benefits of Public Health has made work like this possible. The authors would like to thank the Canadian Nosocomial Infection Surveillance Program (https://health-infobase.canada.ca/cnisp/) for the use of isolates in this validation study. The Public Health Agency of Canada provided funding for the Canadian Nosocomial Infection Surveillance Program. We thank the physicians, epidemiologists, infection control practitioners, and laboratory staff at each participating hospital for their contributions to the study. We gratefully acknowledge the Bioinformatics Core Facility of the National Microbiology Laboratory, Public Health Agency of Canada, for computational infrastructure.

